# Visualizing and integrating linear and graph pangenomes at the Maize Genetics and Genomics Database

**DOI:** 10.64898/2026.08.31.748368

**Authors:** John L. Portwood, Ethalinda K. Cannon, Olivia C. Haley, Laura E. Tibbs-Cortes, Carson M. Andorf, Margaret R. Woodhouse

**Affiliations:** Corn Insects and Crop Genetics Research Unit, US Department of Agriculture - Agricultural Research Service, Ames, IA 50011, USA

## Abstract

Genome visualization has existed for as long as assembled genomes, with an array of tools and approaches to meet researchers’ needs. The genome browser is one of these tools, and it has become a critical analytical resource for geneticists, genome biologists, and breeders. The evolution of genome browser capabilities is ongoing, and the USDA-ARS Maize Genetics and Genomics Database (MaizeGDB) has implemented JBrowse2, the most recent iteration of JBrowse. It includes multiple genome browser and alignment views, demonstrating pangenome visualization capability and permitting sophisticated functional characterization of maize loci. Described here also is MaizeGDB’s new Pangenome Viewer, which integrates pangenome graph views with linear browser visualization to obtain an on-the-fly, interactive visual snapshot of structural variation across a pangenome at user-selected loci, with zoom capabilities and statistical and variant information for each subgraph. Finally, we demonstrate how different MaizeGDB pangenome pipelines complement one another to help guide accurate analyses. This functionality can lead to more precise characterization of loci that confer important agronomic traits, resulting in better outcomes for farmers and the public.

## INTRODUCTION

Genome visualization methods have existed for as long as assembled genomes, with a broad array of tools and approaches available to meet the goals of users. The genome browser is one of these tools, and it has become a critical analytical resource for geneticists, genome biologists, and breeders. The sequencing of the human genome in 2000 prompted the need for a tool to visualize genomic data in a way that was useful to researchers, and the Ensembl genome browser (1) https://www.ensembl.org, hosted by the European Bioinformatics Institute (EBI) (2) https://www.ebi.ac.uk/, was created to support the newly sequenced human genome. The UCSC browser (2, 3)https://genome.ucsc.edu/hostedx by the University of California, Santa Cruz Genomics Institute, was also established around this time. Since then, many other browser platforms have emerged (reviewed in (4)). Some of the most popular browser platforms were created by the Generic Model Organism Database project (GMOD) https://gmod.org/, including GBrowse (4, 5), JBrowse (6), and JBrowse2 (7), which are frequently incorporated into public genome databases.

A typical genome browser is based on the genomic coordinates of a reference genome, against which gene model annotations, variant data, expression data, and other types of evidence are aligned. The browser then aligns these data types along the genomic coordinates, allowing researchers to visualize the locations of these features to identify, for instance, the location of SNPs relative to a gene of interest, or observe the expression patterns of a gene, or assess other functional information. This functionality helps researchers to clone (8-10), characterize (11-13), and otherwise study loci of interest in genetics, genomics, and breeding programs.

The evolution of genome browser capabilities is ongoing. A prime example of browser evolution is the GMOD suite of browsers, beginning with GBrowse, released in 2002. GBrowse was designed to be a web-based application that any user or database could host. Features include the ability to zoom in and out, search loci based on ID or genomic coordinates, and select and organize tracks of interest. GMOD updated their browser platform to JBrowse in 2009. Unlike GBrowse, which relied on server-side rendering, JBrowse relies on client-side rendering, therefore requiring fewer computational resources for the host server. JBrowse’s sidebar menu is also easier to navigate, and the visualization is cleaner and more streamlined. In 2023, GMOD released JBrowse2, which greatly expanded browser functionality by introducing the linear synteny viewer, which allows the comparison of target regions among genome browsers from different accessions or species, including the capacity to visualize whole-genome sequence alignment between two or more genomes, allowing pangenome visualization by selecting multiple genome alignments at once.

Genome browsers, however, are often limited when it comes to viewing pangenome graphs. While pangenomes are collections of genome alignments that allow researchers to analyze the genetic variant relationships across genomes while preserving sequence continuity and structural variation within each genome, a pangenome graph is a single, non-linear entity that represents the genetic diversity of a cohort. A pangenome graph classifies genomic sequences as nodes and their relationships as edges, permitting efficient comparison and analysis of genetic variation within that clade (14). Various pangenome graph pipelines are available, including PGGB (15), which produces an all-to-all alignment of input sequences, graph induction, and progressive normalization; and cactus-Minigraph (16), a reference-based, whole-genome alignment pipeline that uses an iterative sequence-to-graph mapping approach to construct a graph from a set of genomes. A challenge for those who work with pangenome graphs has been finding visualization tools that connect non-linear pangenome graphs to the more familiar linear genome format, particularly by linking pangenome graph visualizations to linear genome browser instances. One tool that has achieved this is PPanG (16, 17), a rice pangenome visualization tool which merges the graph visualization resource SequenceTubeMap (https://github.com/vgteam/sequencetubemap) with JBrowse2.

The USDA-based Maize Genetics and Genomics Database (MaizeGDB - https://www.maizegdb.org) (18) is the community database for maize researchers, providing informatics resources and data curation to support maize genetics, genomics, and breeding research. *Zea mays ssp. mays* (maize, corn) has been the world’s top production grain crop for over a decade (http://faostat.fao.org/). Directed breeding practices (19, 20) have been used to improve maize for uses such as livestock feed (21) and biofuel (21-23). As a model system, maize has been fundamental to the study of transposable element activity (24-26) and genomic complexity (27-29). As of 2026, MaizeGDB hosts 159 genome assemblies across the genus *Zea*, including version 5 of the B73, the representative maize genome; the 25 maize Nested Association Mapping (NAM) founder line genomes (30); 12 founder inbred lines representing heterotic groups from both the US and China (29); and the teosinte wild relatives of domesticated maize from the Pan-Andropogoneae Project (29, 31). MaizeGDB also hosts a number of tools and resources that integrate these genomes and their data sets, including variant, expression, epigenetic, and binding-site data, so that users can more efficiently study the rich variety of genomic and phenomic data within the *Zea* genus.

Genome browsers are a fundamental component of MaizeGDB functionality. In 2010, MaizeGDB presented its first GBrowse instance of the reference B73 genome (32). In 2021, MaizeGDB expanded its capabilities by evolving into a pangenome-centric platform (32, 33) with the hosting of the newly sequenced NAM founder line genomes and further expanded our pangenomic capabilities by introducing the pan-gene center and pre-computed pan-gene relationships across most genomes hosted at MaizeGDB (18). A key step in the expansion was switching from GBrowse to JBrowse, when MaizeGDB introduced JBrowse instances for B73, the representative maize genome, and the 25 NAM founders in 2021. MaizeGDB also generated marker and gene model annotation tracks that allowed users to easily move from one genome’s browser track to another through embedded links within the pop-up boxes of syntenic loci. Since then, the scope of the MaizeGDB genome browsers has expanded substantially to integrate genome-wide comparative, functional, structural, regulatory, and genetic datasets, including syntenome alignments and lifted annotations across grasses, more than 300 geneexpression tracks, over 600 epigenetic and DNA-binding datasets, full-proteome alignments from more than 20 species, AlphaFold and ESMFold protein-structure evidence, multiple alternative gene annotations, maize pangenome and pan-gene datasets, and extensive variant and insertional-mutagenesis resources. This capability resulted in a three-fold increase in browser usage at MaizeGDB (34), and the browsers continue to be one of our most popular features, aiding researchers in cloning (8-10, 35, 36) and characterizing (11-13, 37) loci in maize.

Building upon our browser functionality, we describe MaizeGDB’s upgrade to JBrowse2, https://jbrowse2.maizegdb.org, which permits greatly expanded browser capabilities, such as visualization of whole-genome sequence alignments between two or more genomes, pangenome visualizations, comparisons of RNA expression and syntenic gene model structure across genomes, and other analyses that enhance MaizeGDB’s pangenome-centric platform.

We include tutorials and examples demonstrating how users can study cases such as the impact of indels between syntenic orthologs on expression differences; evaluating gene model annotations; whole-genome sequencing variants; and other phenomena. These examples can serve as a template for other databases to incorporate JBrowse2 into their own platforms. We also introduce MaizeGDB’s Pangenome Viewer, a custom pangenome graph viewer specifically designed for the size, complexity, and unique features of the maize pangenome. This tool generates subgraph pangenome visualizations on the fly from a user-provided gene model identifier or genomic coordinates of the reference genome. The Pangenome Viewer links both reference (currently B73 version 5) and query genomes within the subgraph to their respective browsers and permits users to zoom into a region of interest within the subgraph, then calculate subgraph statistics and analyze variant information. Together, these resources allow MaizeGDB users to explore pangenome visualization and analyses in multiple ways, and we demonstrate use cases that show how different pangenome pipelines can result in complementary outcomes. This functionality can lead to more precise characterization of loci that confer important agronomic traits, resulting in better outcomes for farmers and the public.

## METHODS

### JBrowse2 setup and whole-genome alignment tracks

#### JBrowse2 installation and setup

The web-based version of JBrowse2 was deployed on MaizeGDB infrastructure and is publicly accessible at http://maizegdb.org/genomebrowser. The updated browser hosts 47 maize genome assemblies comprising 6,063 total tracks. All legacy track configurations, customizations, and associated metadata were successfully migrated from MaizeGDB’s JBrowse1 instance. While the initial load time is approximately 15-20 seconds - attributable to processing a comprehensive configuration file encompassing all tracks across every genome - subsequent performance remains robust, even when simultaneously rendering multiple genomes and dozens of tracks.

Beyond native JBrowse2 features, such as single LinearGenomeViews and multi-genome LinearSyntenyViews, we developed custom plugins tailored to maize comparative genomics workflows. For instance, the reference genome assembly (Zm-B73-REFERENCENAM-5.0) incorporates resequencing tracks displaying SNPs identified by (38) relative to other NAM founder lines (e.g., B97, CML103). Selecting a SNP within these tracks launches the standard “Feature Details” pop-up window, augmented by a newly integrated “Pangenome Navigator” section. Clicking this link dynamically initializes a synteny view centered on the selected variant, aligning the B73 reference browser with the corresponding NAM genome browser. In this comparative view, the target SNP is simultaneously highlighted across both windows, enabling immediate visual assessment of local synteny and variant context. Equivalent plugins were engineered for all pangenome-scale tracks in JBrowse2.

#### Alignment track creation

The program AnchorWave v1.2.5 https://github.com/baoxingsong/AnchorWave (39) was used to generate the alignment .paf files for the JBrowse2 linear synteny views and synteny tracks. Pairwise alignment of each of the 44 key browser genomes was performed with the pipeline anchorwave genoAli with ‘-IV true’ against every other genome in MaizeGDB. The resulting .maf files were converted to .paf files using the program wgatools v0.1.0 https://github.com/wjwei-handsome/wgatools (40) in an in-house script. All genomes were downloaded in the summer of 2025. Supplementary Table 1 shows the genome inputs used and other key information.

### Pangenome graph visualization tool

#### Pangenome graph creation

Cactus-Minigraph v9.1.2 (16, 40) was used to generate a graph pangenome from the reference B73 version 5 genome and the 25 Nested Association Mapping (NAM) founder genomes (30). The genomes were downloaded from MaizeGDB (https://download.maizegdb.org/) in December 2025. Cactus was run on the ten maize pseudomolecule chromosomes individually, using default parameters and with the flags ‘--vcf -- gfa --gbz --xg’. Chromosomal .gfa output files were converted to an optimized dynamic genome/graph implementation (odgi (41)) .og file using vg https://github.com/vgteam/vg (42) v1.62.0 convert ‘-f --no-translation’, followed by odgi v0.9.4 build https://github.com/pangenome/odgi. The chromosomal .og and .xg files are used as inputs for the Pangenome Viewer described below.

#### MaizeGDB Pangenome Viewer implementation

The MaizeGDB Pangenome Viewer is implemented as a Flask v3.1.3 (https://flask.palletsprojects.com) web application written in Python 3.14.6 and containerized using Docker for reproducible deployment. The container is built on Ubuntu 22.04 and bundles all required bioinformatics tools, including odgi v0.9.4 and vg v1.62.0, installed via Micromamba from the Bioconda channel (42, 43). The application is served in production using Gunicorn v26.0.0.

#### Pangenome graph extraction and visualization

For each user query, either a B73 version 5 gene model identifier or a set of genomic coordinates, the tool extracts a subgraph of the NAM pangenome from pre-built chromosomal variation graph indexes (.xg files) using ‘vg find’, with fallback to ‘odgi extract’ for regions where ‘vg find’ returns an incomplete result. Extracted subgraphs are converted to .gfa format and built into odgi variation graphs using ‘odgi build’. Gene model annotation paths (CDS and UTR features) are injected into the graph using ‘odgi procbed’ and ‘odgi inject’, and the graph is sorted using ‘odgi sort’ with the Ygs sorting scheme for visual clarity. A static pangenome image is generated using ‘odgi viz’.

An interactive .html visualization of the untangled pangenome graph is generated by a custom .gfa-to-.tsv parsing script (‘gfa_to_untangle_tsv.sh’) that maps each haplotype’s path through the subgraph to both pangenome-local and genomic coordinates, using contig offset information extracted from the chromosomal .xg indexes via ‘vg paths -M’. The .tsv file is rendered as an interactive Plotly (https://plotly.com) figure using R v4.1.2 with ggplot2 v4.0.3 (44) and Plotly v4.12.0 https://plot.lypackages.

Variant calling relative to B73 is performed using ‘vg deconstruct’ on the pre-injection .gfa, with a .gbwt index built from the subgraph using ‘vg gbwt’ to handle path name formats produced by odgi. Summary statistics for each subgraph, including node degree distribution, path depth, and a variant heatmap, are visualized as a four-panel .png using ggplot2.

Code development, debugging, and iterative refinement of the visualization pipeline were assisted by Claude (Anthropic; https://www.anthropic.com), an AI assistant, across multiple development sessions. All code was reviewed and validated by the authors.

#### Static pangenome image generation for gene model pages

The same extraction and visualization pipeline was used to pre-generate static odgi pangenome images for all B73 version 5 gene models. These images are displayed on individual gene model pages on MaizeGDB.

## RESULTS

### JBrowse2 improves pangenomic analyses compared to JBrowse

As of September 2026, MaizeGDB hosts 47 JBrowse instances across a diverse set of maize cultivars and wild relatives. The genomes hosted in these browsers include version 5 of the reference B73 genome and the 25 maize Nested Association Mapping (NAM) founder line genomes (30), 12 founder inbred lines representing heterotic groups from both the US and China (29), and eight teosinte wild relatives of domesticated maize from the Pan-Andropogoneae Project (31). We imported these genomes and their associated browser tracks into JBrowse2. The MaizeGDB JBrowse2 instance can be found here: https://jbrowse2.maizegdb.org/. MaizeGDB also has a browser landing page https://maizegdb.org/genomebrowser where the user can find quick links to access synteny views between any two MaizeGDB JBrowse2 instances for a given genomic coordinate. The landing page includes examples on how to use the JBrowse instance under the “Examples” tab.

One of the key innovations in JBrowse2 is its ability to host whole-genome alignment tracks across multiple genomes (Figure 1). Whole-genome alignments can be generated by a number of different pipelines, provided that the output files are in either .chain or .paf file formats or can be converted to these formats. MaizeGDB performed all-against-all whole-genome alignments across the genomes we host in JBrowse2 using the tool AnchorWave (Methods). We then converted the output .maf files to .paf files to make the alignment tracks for the JBrowse2 browsers (Methods).

**Figure 1.**
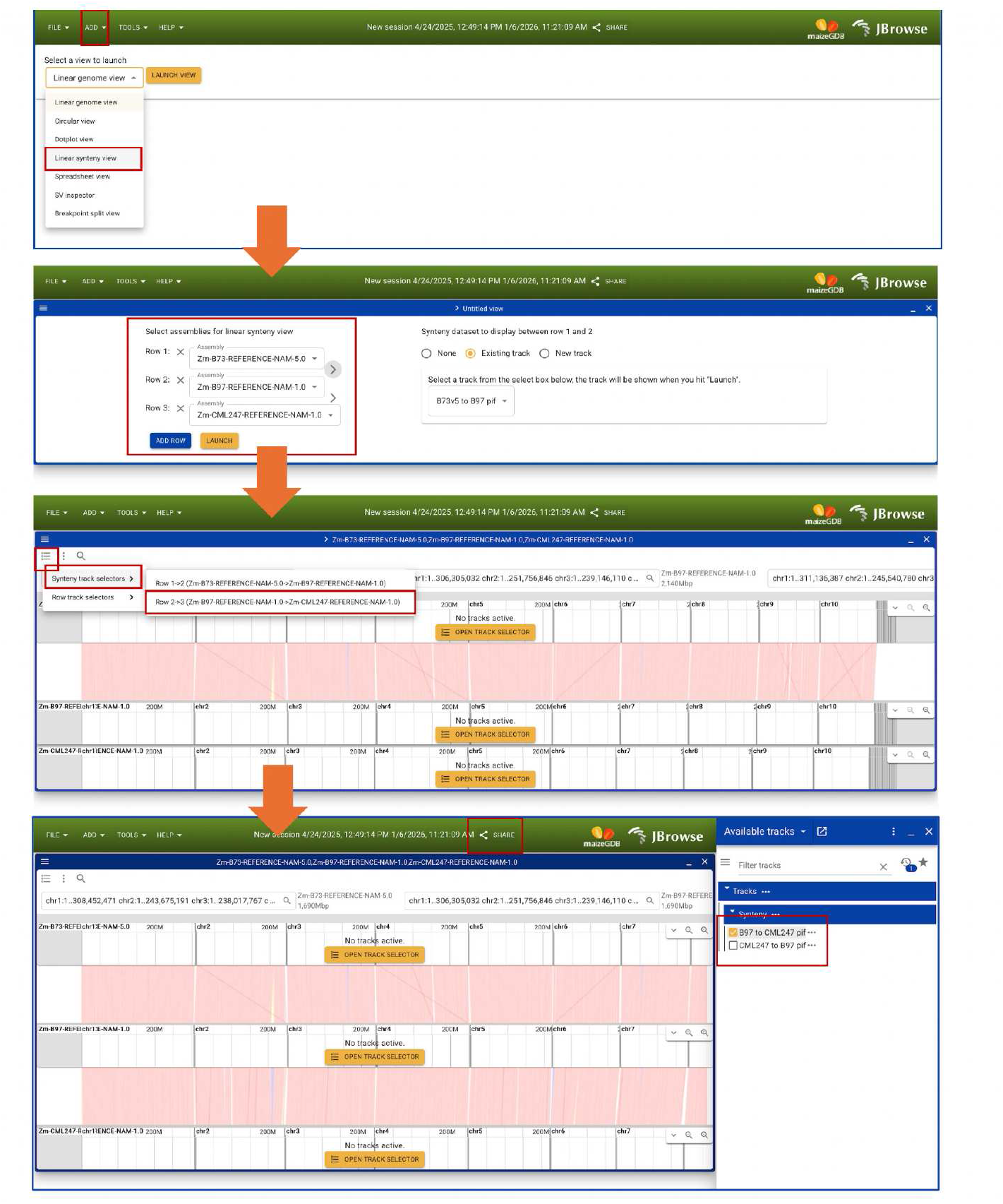
JBrowse2 linear synteny view usage at MaizeGDB https://jbrowse2.maizegdb.org. First, select “Linear synteny view” from the drop-down menu. Then, select the genome assemblies for the linear synteny view (in this example, B73, B97, and CML247); the whole-genome alignment view between the first two genomes (B73 and B97) will be created. Next, go to Synteny track selectors and select the next genomes to view their synteny alignment (B97 and CML247). This will open up a window on the right of the browser to select the alignment track, which will draw the alignment view. The alignment view will then be drawn between B97 and CML247 (right-hand box).

To demonstrate the power and utility of JBrowse2, we show in Figure 2 the difference between using JBrowse to compare structural differences in syntenic gene models across genomes vs using JBrowse2. Figure 2A shows a JBrowse instance of gene model Zm00001eb017450 in the reference genome B73 (top) and the syntenic orthologs of two other genomes from the NAM founder assemblies, M162W (middle) and CML69 (bottom), lifted to the B73 genomic coordinates. The reference B73 gene model Zm00001eb017450 and its B97 syntenic ortholog contain two exons that are missing in the CML69 gene model Zm00020ab017810 (rectangle). Clicking on the CML69 gene model Zm00020ab017810 opens up a pop-up box with a URL to the CML69 JBrowse instance to view the gene model Zm00020ab017810 in the CML69 browser. We can confirm that the CML69 gene model Zm00020ab017810 is indeed missing those two exons; the RNA-seq bigwig file tracks below it show no expression where those exons would be if the exons had been excluded due to an annotation error.

**Figure 2.**
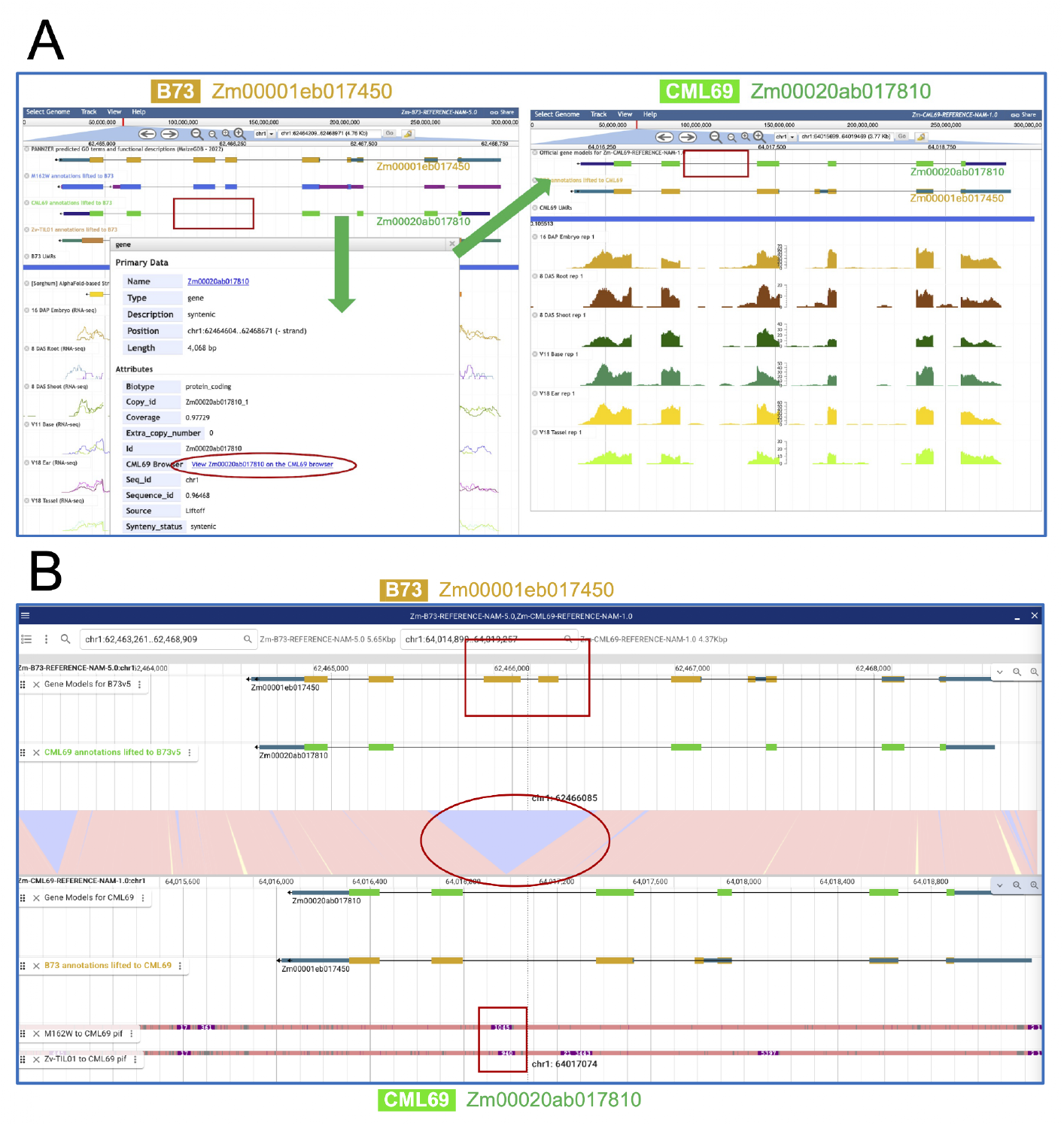
JBrowse2 improves pangenomic analyses compared to JBrowse. A) JBrowse instance of gene model Zm00001eb017450 in the reference genome B73 (top) and syntenic orthologs of NAM founder genomes B97 (middle) and CML69 (bottom), lifted to B73 genomic coordinates. The B73 gene model Zm00001eb017450 and the M162W ortholog contain two exons that are missing in the CML69 gene model Zm00020ab017810 (rectangle). When viewing the CML69 gene model Zm00020ab017810 in the CML69 browser, we see it is indeed missing as two RNA-seq bigwig file tracks (selected from menu, not shown) show no expression beneath those exons. B) The JBrowse2 linear synteny view of the B73 gene model Zm00001eb017450 (top) and the CML69 gene model Zm00020ab017810 (bottom). The exons in the B73 gene model (rectangle) are missing in the CML69 syntenic ortholog; the blue space (circled) indicates DNA sequence in B73 that has been deleted in CML69. Alignment tracks to B97 and TIL01 in the CML69 browser show that this sequence (rectangle) that has been deleted in CML69 is present in those two assemblies.

Compare Figure 2A with Figure 2B, which shows the same gene models, the reference B73 gene model Zm00001eb017450 (top) and the CML69 gene model Zm00020ab017810 (bottom), in a linear synteny alignment view between the two genomes in JBrowse2. We can clearly see that the exons in the B73 gene model (rectangle) are missing in the CML69 syntenic ortholog by observing the alignment visualization track between the two genomes. The blue space (circled) indicates DNA sequence in B73 that has been deleted in CML69. Furthermore, we can select alignments (using the same .paf files that create the linear synteny views) between CML69 and B97 and TIL01 as tracks in the CML69 browser to show that this deleted sequence (rectangle) is also present in the syntenic regions in those two assemblies. This illustrates how JBrowse2 can more simply and clearly show structural variation among syntenic gene models with alignment support.

### Studying the functional effects of post-polyploid fractionation in JBrowse2

A more complex example is featured in Figure 3 comparing a JBrowse2 linear synteny view of the B73 gene model Zm00001eb051200 and its NAM founder genome NC358 syntenic ortholog Zm00037ab051600. The top panel (Figure 3A) shows the JBrowse2 B73 genome browser, with the B73 gene model track highlighted, including Zm00001eb051200. Below it is the lifted NC358 annotation track featuring Zm00037ab051600 (arrow), followed by the lifted Sorghum syntenic gene model annotation Sobic.001G162700, then a G4-quadruplex track, then the lifted syntenic gene model annotations of the NAM founder genomes CML130 and CML247, respectively. These tracks are followed by a gene model track by the ReelGene machine-learning pipeline (45), which scores annotation quality; the exons in light blue are high-confidence exon annotations. Below this track is the Peptide Atlas track that uses The Trans-Proteomic Pipeline to analyze maize tandem mass spectrometry to identify gene models that generate peptides (46) https://peptideatlas.org/builds/maize/. This is followed by a track of unmethylated regions (UMRs), predictive of gene function; predicted transposable elements; and RNA-seq from anther tissue (10.1126/science.abg5289). Below this is the alignment (Figure 3B) between B73 and NC358 at that locus. Then, the bottom panel is the Zm00037ab051600 gene model (arrow) in its own NC358 genome browser (Figure 3C) (note that Zm00037ab051600 may be misannotated and missing its 3’ exon when compared to the lifted over B73 syntenic ortholog below).

**Figure 3.**
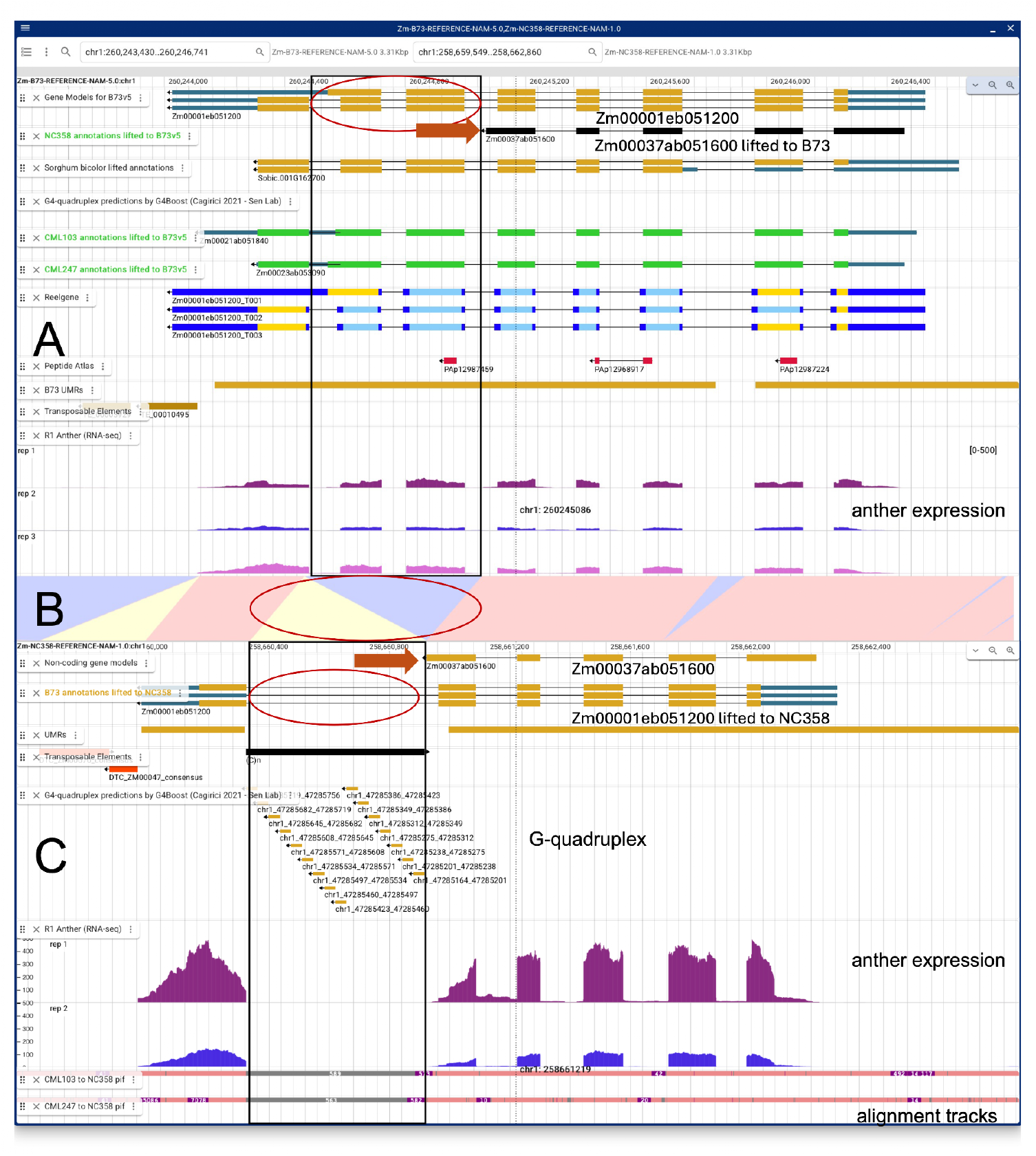
Studying functional effects of post-polyploid fractionation in JBrowse2 featuring a JBrowse2 linear synteny view of the B73 gene model Zm00001eb051200 and its NAM founder genome NC358 genome syntenic ortholog Zm00037ab051600. A) B73 genome browser. Top track: B73 gene model track, featuring Zm00001eb051200. Below is the lifted NC358 annotation track featuring Zm00037ab051600 (arrow); lifted Sorghum syntenic gene model annotation Sobic.001G162700; lifted gene model annotations of NAM genomes CML130 and CML247; ReelGene annotation quality scores (blue exons=high-confidence exon annotations; Peptide Atlas track; unmethylated regions (UMRs), predictive of gene function; predicted transposable elements; RNA-seq of anther tissue. B) Alignment between B73 and NC358. C) Zm00037ab051600 gene model (arrow) in its own NC358 genome browser. Two of the exons shared across B73, Sorghum, CML103, and CML247 (circled, black rectangles) are missing in NC358. The NC358 gene has a predicted G4-quadruplex repeat element inserted downstream from this deletion. While the NC358 genome model Zm00037ab051600 has undergone deletion, it is still expressed, and expressed more highly in anthers than in B73.

We can observe from this example that two of the exons shared across B73, Sorghum, CML103, and CML247 (circled, black rectangles) are missing in NC358. Furthermore, NC358 has a repeat element inserted just downstream from this deletion, which turns out to be a predicted G4-quadruplex (47), a type of non-B DNA structure (Fig. 3C). It has been shown that G4 structures can impede DNA replication and promote recombination and genome rearrangements, including deletions (48, 49), and G4-forming sequences have been identified within plant retrotransposons and proposed as sites of post-insertional genome rearrangement (50). Although these data do not establish that the G4 contributed to the NC358 deletion, they illustrate how integrated genome-browser tracks can reveal candidate sequence features associated with structural genome evolution. Also of note, this NC358 locus was previously observed to have undergone fractionation, or partial gene deletion, relative to its homeolog when compared to its Sorghum ortholog (30). A careful study of these data demonstrates that while the NC358 genome model Zm00037ab051600 has undergone deletion, it is still expressed, and expressed more highly in anthers than in B73, as observed by the example anther RNA-seq bigwig tracks below the gene models in each genome. Interestingly, this gene model was originally predicted to be noncoding. This example shows how researchers can use the MaizeGDB JBrowse2 instances to study genome evolution at loci of interest and its impact on gene function.

### Using JBrowse2 to study variant effect functional differences between genomes

The MaizeGDB tool SNPversity 2.0 https://wgs.maizegdb.org/ is a web-based tool to visualize and analyze maize variant data (30, 51). In Figure 4, we show a stop-loss predicted high-impact variant In SNPversity for the gene model Zm00001eb407010 (Figure 4A). We see that this stop-loss variant segregates across the NAM founder population. The tool includes links to the B73 JBrowse instance highlighting each variant, as shown in Figure 4B. In the B73 JBrowse instance, we can compare the B73 gene model annotation to gene model annotations lifted from a subset of the NAM founder genomes to B73. We can see that B73, B97, and Oh43, all which are absent the stop-loss variant, have a truncated 3’ region of their gene models compared to CML277, CML333, and Tzi8, which do have the stop-loss variant.

**Figure 4.**
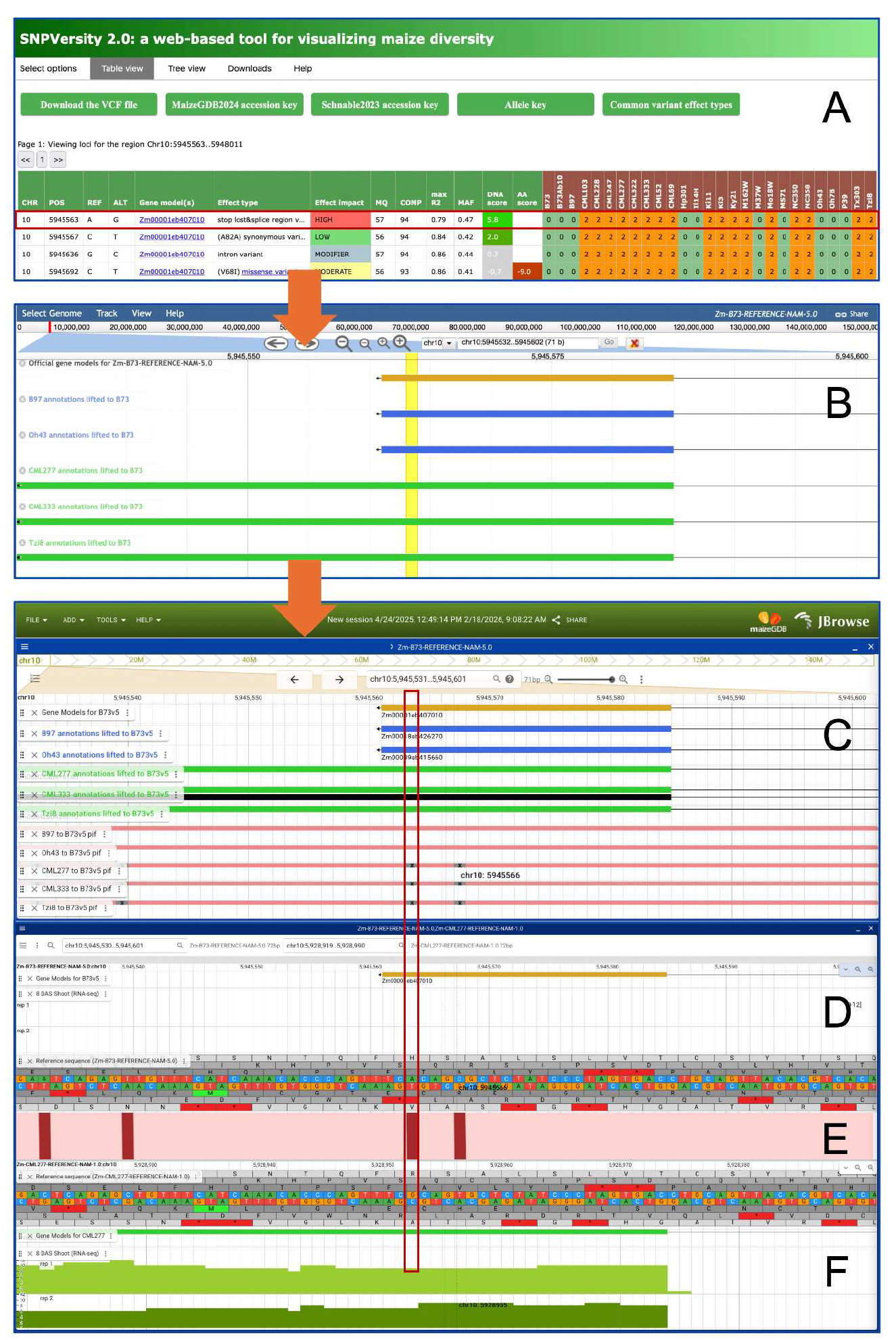
Using JBrowse2 to study variant effect functional differences between genomes. A) A stop-loss predicted high-impact variant In SNPversity for the gene model Zm00001eb407010. B) The B73 JBrowse instance highlighting the stop-loss variant. We can compare the B73 gene model annotation to annotations lifted from NAM genomes B73, B97, and Oh43, all of which are absent the stop-loss variant and have a truncated 3’ region of their gene models, and CML277, CML333, and Tzi8, which do have the stop-loss variant. C) JBrowse2 single linear B73 instance of the same lifted gene models with the same variant region highlighted but with alignment tracks for the genomes of the lifted gene models, we see the variant locus for the genomes that have the stop-loss variant (CML227, CML333, and Tzi8). D) The linear synteny view between B73 and E) its alignment window to F) CML277, one of the genomes missing the stop-loss variant. The variant base-pair change is highlighted. Shoot RNA-seq tracks demonstrate that the gene locus is transcribed.

Below this JBrowse panel is a JBrowse2 single linear B73 instance of the same lifted gene models with the same variant region highlighted (Figure 4C). Here, however, we also have alignment tracks for the genomes of the represented lifted gene models, and it’s easy to see the variant locus for the genomes that have the stop-loss variant (CML227, CML333, and Tzi8). Below that panel is a linear synteny view between B73 (Figure 4D) and its alignment (Figure 4E) to CML277 (Figure 4F), one of the genomes missing the stop-loss variant. Once again, the actual variant and the base-pair change responsible is highlighted. In the synteny alignment, we can see the base-pair difference between B73 and CML277, which codes for the stop-loss variant, as predicted by SNPversity. In this way, JBrowse2 allows users to compare variant data between two or more genomes and observe multiple data types at once.

### Pangenome graph visualization tool

Sometimes there are cases where a user would prefer a quick snapshot of the structural variability within a given locus. The alignment tracks on the JBrowse2 instances allow users to easily compare two or more genomes at a given locus. However, a user who would like to observe the alignments of a genome cohort must manually select multiple alignment tracks and modify track height and other parameters in order to view the whole alignment landscape at once. A more straightforward way to view pangenomic alignments at a given locus is to generate images from a graph pangenome on demand. Here, we introduce MaizeGDB’s graph Pangenome Viewer https://pangenome-viewer.maizegdb.org/, where a user can select the pangenome of their choice, enter either their reference B73 version 5 gene model of interest or the genomic coordinates of a B73 locus, then view a static pangenome subgraph and an interactive untangled subgraph at the selected locus (Figure 5). Currently, this tool hosts a cactus-Minigraph pangenome (Methods) of the 25 NAM founder assemblies and the reference B73 v5 genome (30).

**Figure 5.**
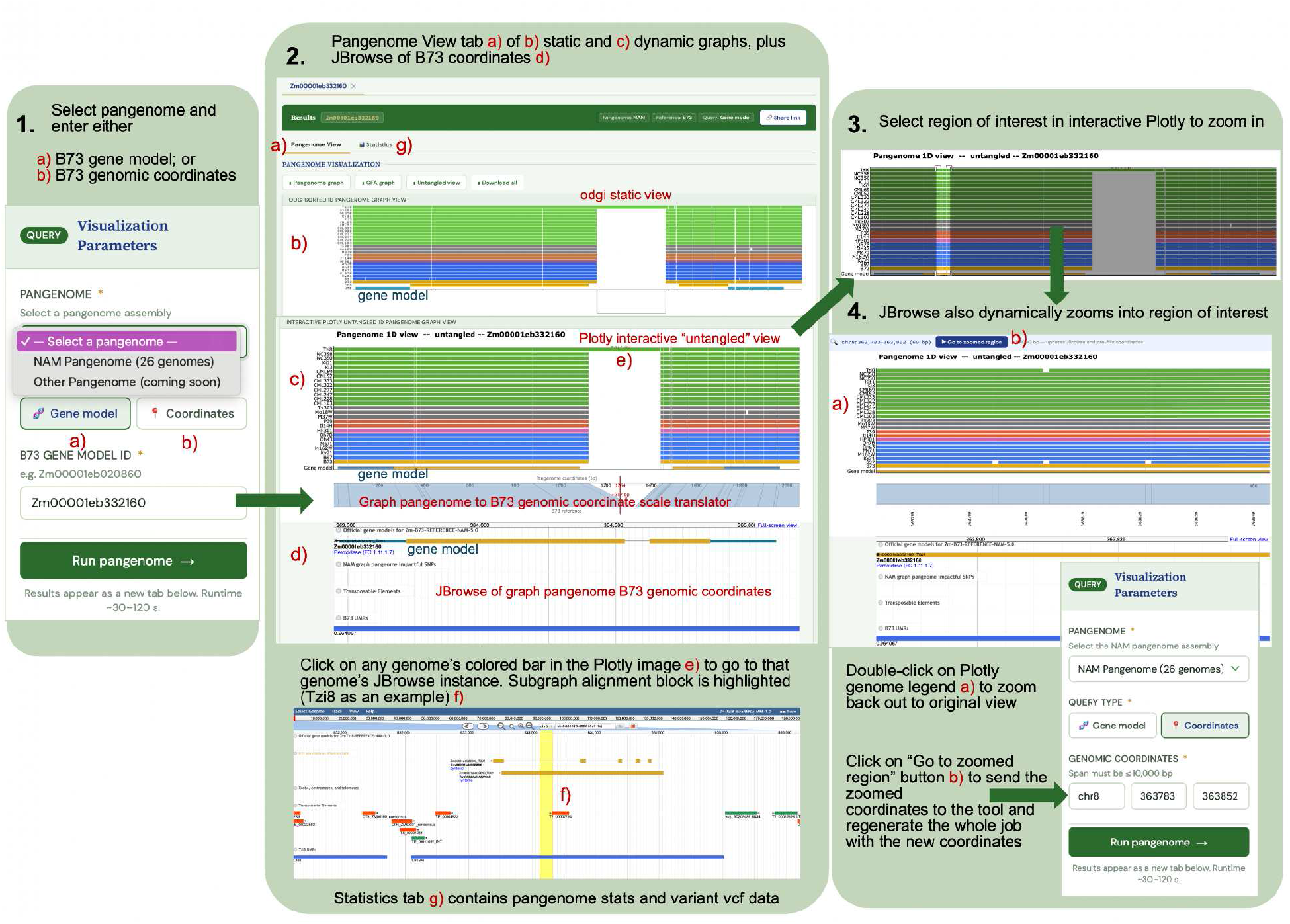
the quick-start image on the Pangenome Viewer homepage. 1: Users select a pangenome and enter either a B73 version 5 gene model or genomic coordinates. 2: The Pangenome Viewer output. Shown is the Pangenome view tab, with the static odgi subgraph at the top, the interactive Plotly untangled subgraph in the middle, and the B73 version 5 JBrowse instance below with the same genomic coordinates as the B73 genomic subgraph coordinates. For both subgraph views, the CDS regions are in gold and the UTRs are in aqua blue. Below is the insertion region in Tzi8 shown in Tzi8’s home JBrowse instance, highlighted in yellow. 3: To zoom in, select region of interest. JBrowse view will also dynamically zoom in. To re-run job so that the odgi static view and the statistics are re-calculated, click on the “Go to zoomed region” button and click “Run pangenome” again.

The graph Pangenome Viewer tool has a Flask (https://flask.palletsprojects.com/en/stable/) back-end and relies on Python (https://www.python.org/), R (https://www.R-project.org/), Plotly (https://plotly.com), ggplot2 (30, 44), odgi (41), and vg (42) (https://github.com/vgteam/vg) to generate the images (Methods). The tool has two tabs: the Pangenome View tab, and the Statistics tab (Figure 5). The Pangenome View tab shows the graph pangenome two ways: a static odgi view of the selected locus organized by haplotype group (i.e. for the NAM pangenome, green=tropical, blue=nonstiff-stalk, gray=mixed, pink=popcorn, orange=sweet corn, yellow=stiff-stalk), and a similar dynamic Plotly view of the untangled graph pangenome. Both views also show the merged UTR and CDS features within the locus, if present.

Below the Plotly view is the MaizeGDB B73 version 5 JBrowse instance for the B73 genomic regions shown in the Plotly view. The JBrowse view shows the gene model annotation tracks, SNPEff-annotated impactful SNPs derived from the cactus-Minigraph pangenome, the transposable element track so that users can see whether or not complex regions in the graph overlap repeat regions in the reference, and an unmethylated region (UMR) track, indicative of coding regions. Observable in Figure 5 is that the gene model in JBrowse lines up precisely with the Plotly graph gene model. Between the Plotly view and the JBrowse instance is a scale that shows the local pangenome coordinates at the top, with the bottom representing the B73 genomic coordinates that match the coordinates in the JBrowse instance below it. This functionality connects the graph pangenome to the linear reference genome for direct analyses of gene models or other regions of interest.

In Figure 5, a gap exists between the pangenomic coordinates and the B73 coordinates (circled), highlighting the insertion in the genome Tzi8 relative to B73. If a user wants to take a closer look at any part of the Plotly image, they can use their mouse to select a region in the graph, and the graph will auto-zoom to the selected region. The JBrowse coordinates will also automatically re-load to the zoomed region. To return to the original view, the user only has to double-click on the graph image’s genome legend again; the JBrowse coordinates will also revert to the original coordinates. Note that the static odgi view remains the same. If a user wants the static odgi view to zoom in as well, the user simply clicks on the “Go to zoomed region” button above the Plotly graph image, and the tool will load the zoomed-in coordinates to the Visualization parameters on the left and re-run the job. The odgi static image png and the dynamic Plotly html are both downloadable, as is the .gfa file for the selected locus.

To see what a non-B73 genome in the subgraph looks like in its native JBrowse instance, just click on one of the subgraph’s alignment blocks in the alignment bar for that genome and it will take the user to the JBrowse instance within the selected alignment block. In that genome’s JBrowse instance, its gene model, structural features, transposable element, and UMR tracks will open automatically, along with the lifted over B73 gene models for easy comparison.

A URL for each job can be generated by clicking on the “Share link” button at the top right of the page. These links can also be pre-built with a user’s gene model or coordinates of interest. That way, entering the URL in the browser will automatically run the job without having to enter the information into the Visualization Parameters box. A static odgi view for each B73 version 5 gene model input is viewable for the gene model pages of each B73 gene model in MaizeGDB.

The Statistics tab includes a number of statistics that users of graph pangenomes might find useful, such as the number of nodes, the number of edges, the number of haplotype (i.e. genome) paths, as well as bar charts for mean node degree for each genome and mean path depth. Below this is a visualization of variants in each genome within the selected locus, generated by vg deconstruct. All images as well as the vcf file are downloadable. Note that if a user re-runs the entire job on a zoomed-in region using the “Go to zoomed region” button in the Pangenome View tab, these images and statistics will be re-run as well for the zoomed locus. More examples and a FAQ can be found at the tool’s URL.

### When pangenomes disagree

Different pangenome formats are generated using distinct methods. For instance, the MaizeGDB pan-gene data set of the genomes at MaizeGDB, created for MaizeGDB’s pan-gene data center https://maizegdb.org/pan_gene_center/pan_gene (18, 42), was generated by the tool Pandagma (52), which uses a CDS alignment method followed by DagChainer (53) collinearity searches to identify syntenic pan-gene relationships across a clade. By contrast, the JBrowse2 alignment visualization files were generated from whole-genome alignment outputs of the Minimap-based AnchorWave tool https://github.com/baoxingsong/AnchorWave (39, 53), which uses shared locus anchors to identify syntenic alignment blocks, and collectively, these paired alignments can serve as a pan-genome for JBrowse2 visualization. Meanwhile, the graph Pangenome Viewer tool uses cactus-Minigraph, which uses a minimap2-based iterative sequence-to-graph mapping approach to construct a graph from a set of genomes. In most cases, the three different pangenome approaches tend to agree with one another. For example, if we compare the output from the Pangenome Viewer for gene model Zm00001 eb332160 (Figure 6A) to the JBrowse2 (Figure 6B) and Pandagma (Figure 6C) outputs, we find that the three pangenomes show that this locus is retained across all NAM founders, and that the structural variation patterns seen in both the Pangenome Viewer and in JBrowse2 tend to agree.

**Figure 6.**
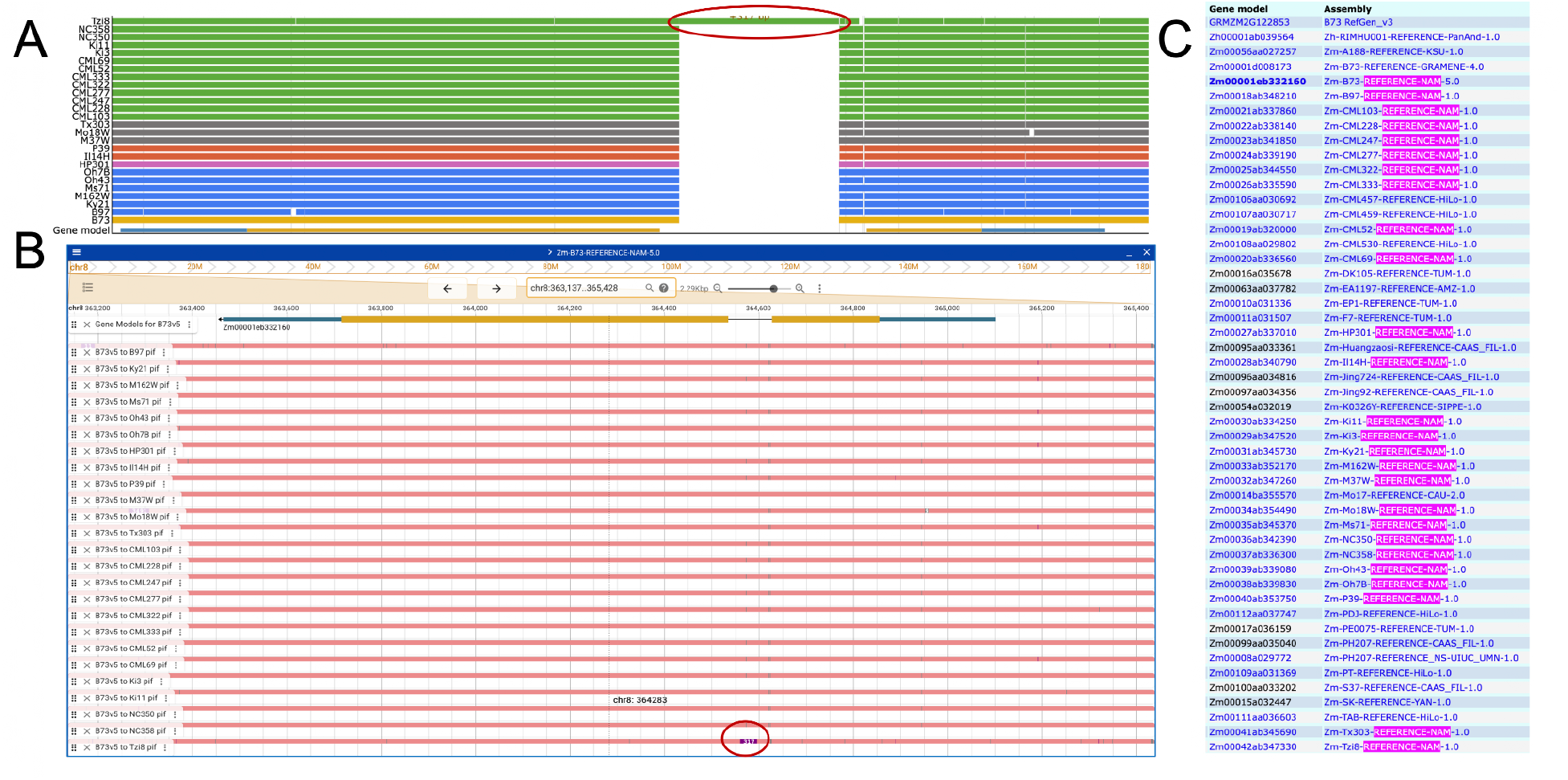
Different pangenome outputs for the B73 gene model Zm00001eb332160 from A) the Pangenome Viewer; B) JBrowse2; and C) Pandagma. The red circles in A and B show the insertion in the. Tzi8 genome. The pink highlighting in Care all the NAM founder genomes represented in the pan gene instance. All three pangenomes show that this gene model is retained across the NAM founder genomes.

However, these three methods are complementary to one another, and sometimes one method will identify pan-genomic relationships that the others do not. For example, the B73 gene model Zm00001eb332410, a MYS-related-transcription factor, has at least a partial alignment for every NAM founder genome in the Pangenome Viewer (though the 3’ end of the gene appears to be missing in the majority of NAM genomes) (Figure 7A). Yet eight of the 25 NAM founder genomes are missing a member in the MaizeGDB Pandagma pan-gene for Zm00001eb332410 (Figure 7C). If we investigate this gene model in JBrowse2 (Figure 7B), we find that the JBrowse2 view more closely matches what is observed in the Pangenome Viewer rather than Pandagma. However, if we click on an alignment block in the Pangenome view associated with a genome that’s missing a representative in Pandagma to view that alignment block in its native JBrowse instance-in this example B97 (Figure 7A, circled)--we see that there is no gene model annotation at that location in B97 for Pandagma to align. We can then conclude that the omission in Pandagma was not due to an issue with the Pandagma pipeline, but rather because of the absence of a gene model in that region within that genome.

**Figure 7.**
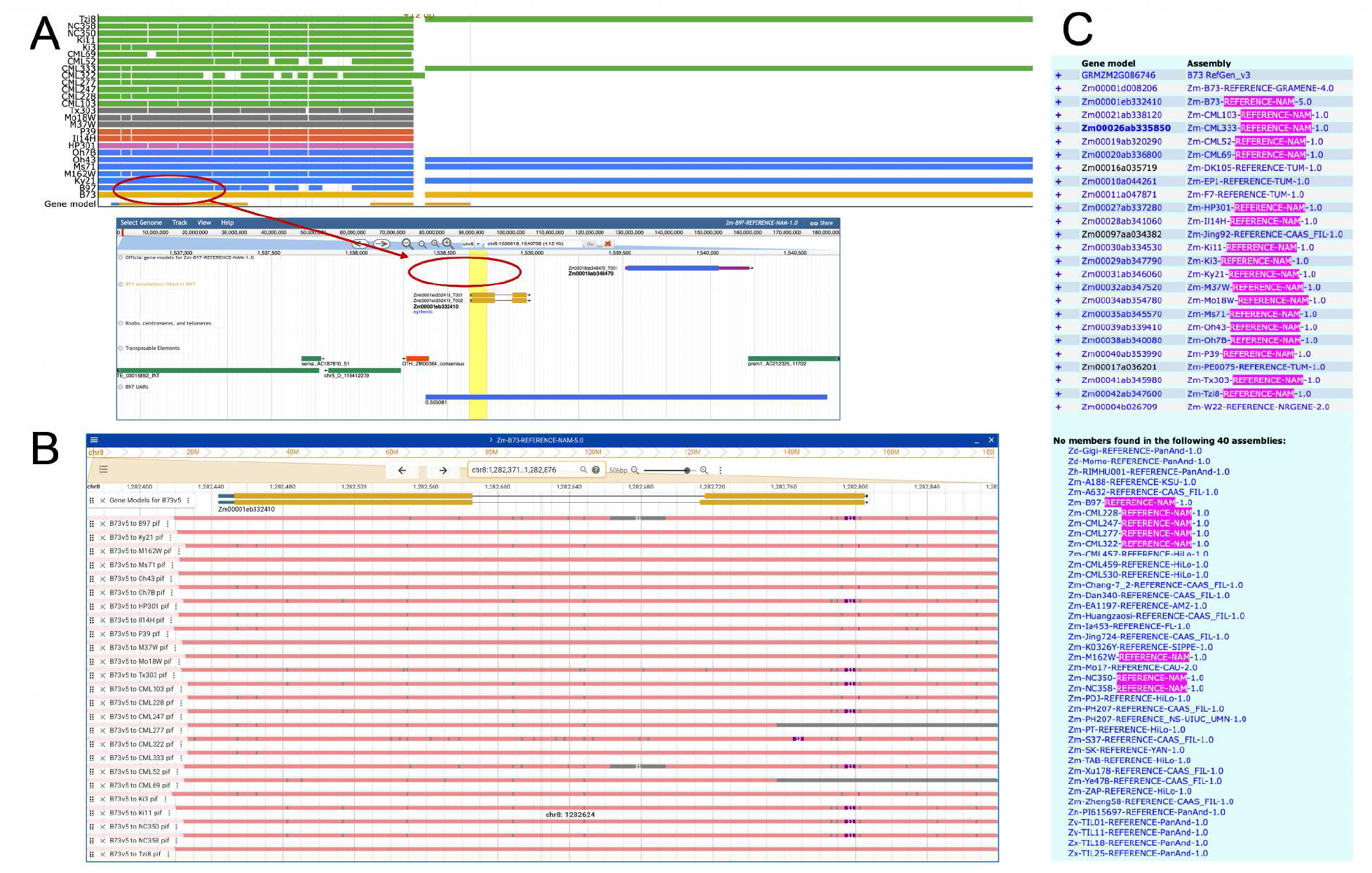
Different pangenome outputs for the B73 gene model Zm00001eb332410 from A) the Pangenome Viewer; B) JBrowse2; and C) Pandagma. Zm00001eb332410 has at least a partial alignment for every NAM founder genome in the Pangenome Viewer, though the 3’ end of the gene appears to be missing in the majority of NAM genomes. But eight of the 25 NAM founder genomes are missing a member in the MaizeGDB Pandagma pan-gene (C). The JBrowse2 view more closely matches what is observed in the Pangenome Viewer than Pandagma. But clicking on an alignment block in the Pangenome view associated B97, a genome missing in the Pandagma pan-gene, to view that alignment block in its native JBrowse instance, we see no gene model annotation at that location in B97 for Pandagma to align.

Pipelines like AnchorWave and cactus-Mingraph are not strictly reliant on gene model annotations to identify shared sequence, and can therefore find sequence similarity where gene models are missing or are incorrectly annotated. Conversely, complex regions, especially those with large indels or repeat regions, can be difficult for graph pipelines to navigate through, whereas a program like Pandagma, which relies solely on gene model alignment and collinearity, can call pan-gene relationships without respect to nearby repeat or indel regions. The following example for the gene Zm00001 eb244620 (Figure 8) shows where Pandagma (Figure 8C) was able to capture almost the full pan-gene alignment across the NAM founders except for M37W, while the majority of NAM founder genomes are not represented in the Pangenome Viewer (Figure 8A). If we take a look at this gene model (circled) in JBrowse2 (Figure 8B), we can see a great deal of complexity in this region (rectangles) relative to B73. We can infer that cactus-Minigraph had trouble aligning through this region and therefore was not able to find paths through many of the genomes.

**Figure 8.**
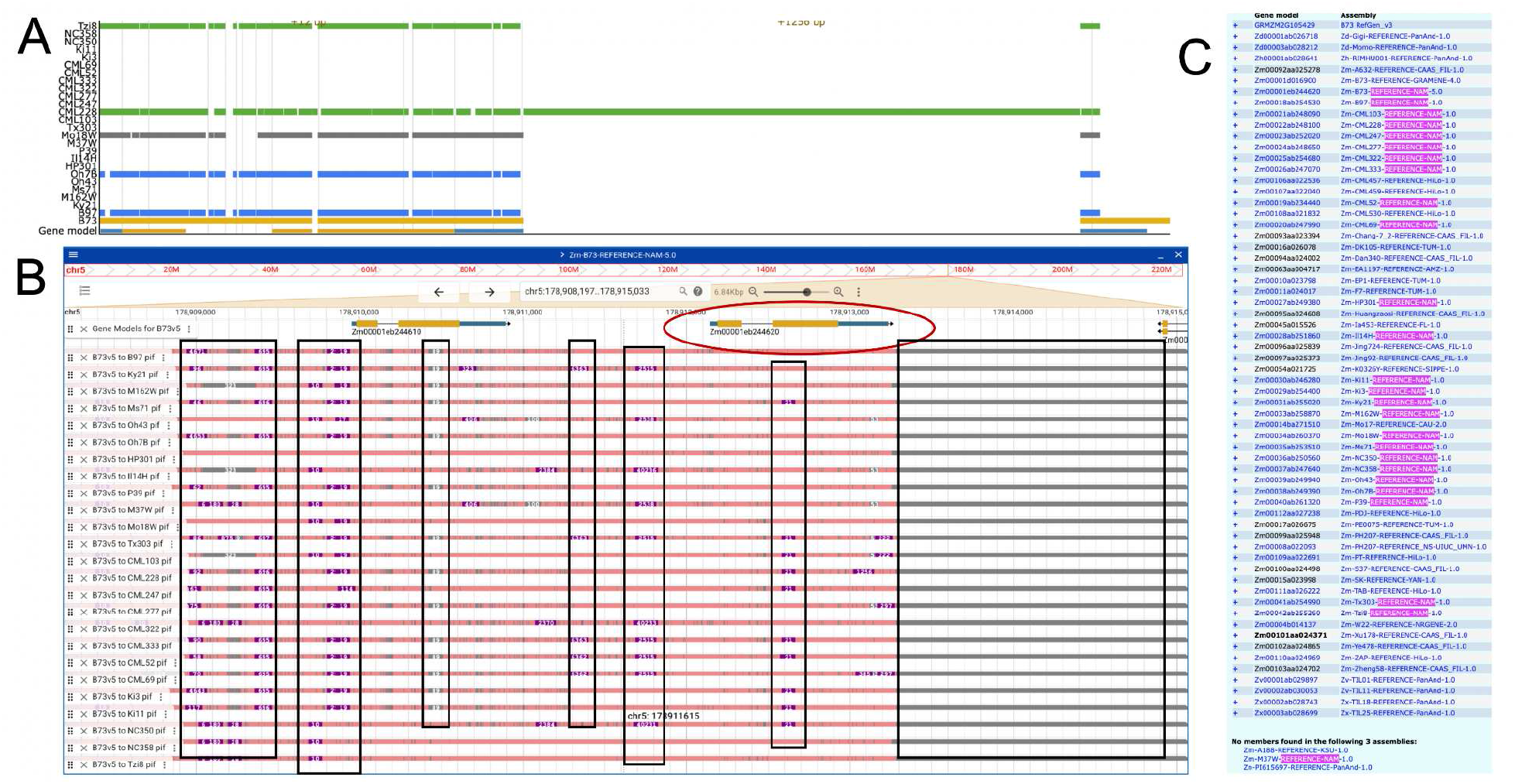
Different pangenome outputs for the B73 gene model Zm00001eb244620 from A) the Pangenome Viewer; B) JBrowse2 (circled); and C) Pandagma. Here, Pandagma (C) was able to capture almost the full pan-gene alignment across the NAM founders except for M37W, while most NAM founder genomes are not represented in the Pangenome Viewer (A). The JBrowse2 view (B) shows significant complexity in this region (rectangles) compared to B73. We can infer that cactus-Minigraph had trouble aligning through this region and therefore was not able to find paths through many of the genomes.

The above examples demonstrate that a single pan-genome pipeline is not sufficient to represent a pan-gene locus in all cases. Structural variation, the absence or misannotation of gene models, and other factors can cause pan-gene loci to be missed in one pipeline but captured in another. It is for this reason that MaizeGDB offers multiple ways to visualize and analyze pangenomes across maize.

## CONCLUSION

Pangenome visualization is constantly evolving and improving along with emerging genomic and pangenomic data and methods. MaizeGDB has implemented new tools and resources from JBrowse2 to the MaizeGDB Pangenome Viewer to keep apace with advancements in genome biology and pangenomics. Here we demonstrated our implementation of JBrowse2 and its usefulness when doing whole-genome comparisons across a cohort of genomes, when assessing differences in gene models across genomes, and when doing deep functional analyses of target genes. We also introduce our new Pangenome Viewer tool, which adds another layer of pangenome visualization for maize users. Finally, we compare and contrast the ways in which different genomic and pangenomic pipelines report the genomic depth of a pan-gene locus, and how multiple ways of analyzing these data can lead to more precise characterization of important loci that confer important agronomic traits, resulting in better outcomes for farmers and the public.

## DATA AVAILABILITY

The MaizeGDB JBrowse2 instance can be found here: https://jbrowse2.maizegdb.org/. MaizeGDB also has a browser landing page https://maizegdb.org/genomebrowser where the user can find quick links to access synteny views between any two MaizeGDB JBrowse2 instances for a given genomic coordinate, or enter a B73 gene model to view the NAM pangenome track views as shown in figures 6-8. The landing page includes examples on how to use the JBrowse2 instance under the “Examples” tab.

The Pangenome Viewer can be accessed at https://pangenome-viewer.maizegdb.org/. The GitHub repository for the Pangenome Viewer can be found at https://github.com/Maize-Genetics-and-Genomics-Database/pangenome_viewer.

## SUPPLEMENTARY DATA STATEMENT

Supplementary Data are available at NAR Online.

## AUTHOR CONTRIBUTIONS STATEMENT

Conceptualization: MRW, JLP. Data curation: JLP, CMA, LET-K, EKC, OCH, MRW. Investigation: MRW, JLP. Methodology: JLP, MRW. Software: MRW, JLP. Visualization: JLP, MRW. Writing - original draft: MRW, JLP. Writing - review & editing: CMA, LET-K, EKC, OCH.

## REFERENCES

1. Dyer, S.C., Austine-Orimoloye, O., Azov, A.G., Barba, M., Barnes, I., Barrera-Enriquez, V.P., Becker, A., Bennett, R., Beracochea, M., Berry, A., et al. (2025) Ensembl 2025. Nucleic Acids Res, 53, D948–D957.

2. Thakur, M., Bosc, N., Brooksbank, C., Ernst, C., Freeberg, M.A., Gurwitz, K.T., Hermjakob, H., Hulcoop, D.G., Martin, M.J., McDonagh, E.M., et al. (2026) EMBL’s European Bioinformatics Institute (EMBL-EBI) in 2025. Nucleic Acids Res, 54, D10–D19.

3. Casper, J., Speir, M.L., Raney, B.J., Perez, G., Nassar, L.R., Lee, C.M., Hinrichs, A.S., Gonzalez, J.N., Fischer, C., Diekhans, M., et al. (2026) The UCSC Genome Browser database: 2026 update. Nucleic Acids Res, 54, D1331–D1335.

4. Wang, J., Kong, L., Gao, G. and Luo, J. (2013) A brief introduction to web-based genome browsers. Brief Bioinform, 14, 131–143.

5. Stein, L.D., Mungall, C., Shu, S., Caudy, M., Mangone, M., Day, A., Nickerson, E., Stajich, J.E., Harris, T.W., Arva, A., et al. (2002) The generic genome browser: a building block for a model organism system database. Genome Res, 12, 1599–1610.

6. Skinner, M.E., Uzilov, A.V., Stein, L.D., Mungall, C.J. and Holmes, I.H. (2009) JBrowse: a next-generation genome browser. Genome Res, 19, 1630–1638.

7. Diesh, C., Stevens, G.J., Xie, P., De Jesus Martinez, T., Hershberg, E.A., Leung, A., Guo, E., Dider, S., Zhang, J., Bridge, C., et al. (2023) JBrowse 2: a modular genome browser with views of synteny and structural variation. Genome Biol, 24, 74.

8. Cao, Y., Zeng, H., Ku, L., Ren, Z., Han, Y., Su, H., Dou, D., Liu, H., Dong, Y., Zhu, F., et al. (2020) ZmIBH1-1 regulates plant architecture in maize. J Exp Bot, 71, 2943–2955.

9. Forestan, C., Farinati, S. and Varotto, S. (2012) The Maize PIN Gene Family of Auxin Transporters. Front Plant Sci, 3, 16.

10. Ozcan, K.E. and Monroe, J.D. (2023) Maize -amylase7 encodes 2 proteins using alternative transcriptional start sites: Nuclear BAM7 and plastidic BAM2. Plant Physiol, 192, 2871–2882.

11. Barnes, A.C., Rodrfguez-Zapata, F., Juarez-Nufiez, K.A., Gates, D.J., Janzen, G.M., Kur, A., Wang, L., Jensen, S.E., Estevez-Palmas, J.M., Crow, T.M., et al. (2022) An adaptive teosinte introgression modulates phosphatidylcholine levels and is associated with maize flowering time. Proc Natl Acad Sci U S A, 119, e2100036119.

12. Trentin, H.U., Krause, M.D., Zunjare, R.U., Almeida, V.C., Peterlini, E., Rotarenco, V., Frei, U.K., Beavis, W.D. and Lubberstedt, T. (2023) Genetic basis of maize maternal haploid induction beyond and. Front Plant Sci, 14, 1218042.

13. Agostini, R.B., Ariel, F., Rius, S.P., Vargas, W.A. and Campos-Bermudez, V.A. (2023) Trichoderma root colonization in maize triggers epigenetic changes in genes related to the jasmonic and salicylic acid pathways that prime defenses against Colletotrichum graminicola leaf infection. J Exp Bot, 74, 2016–2028.

14. Kopalli, V., Arslan, K., Morales-Dfaz, N., Zanini, S.F. and Golicz, A.A. (2025) Toward a standardized framework for pangenome graph evaluation: assessing crop plant pangenome variation graph construction from multiple assemblies. Gigascience, 14.

15. Garrison, E., Guarracino, A., Heumos, S., Villani, F., Bao, Z., Tattini, L., Hagmann, J., Vorbrugg, S., Marco-Sola, S., Kubica, C., et al. (2024) Building pangenome graphs. Nat Methods, 21, 2008–2012.

16. Hickey, G., Monlong, J., Ebler, J., Novak, A.M., Eizenga, J.M., Gao, Y., Human Pangenome Reference Consortium, Marschall, T., Li, H. and Paten, B. (2024) Pangenome graph construction from genome alignments with Minigraph-Cactus. Nat Biotechnol, 42, 663–673.

17. Liu, M., Zhang, F., Lu, H., Xue, H., Dong, X., Li, Z., Xu, J., Wang, W. and Wei, C. (2024) PPanG: a precision pangenome browser enabling nucleotide-level analysis of genomic variations in individual genomes and their graph-based pangenome. BMC Genomics, 25, 405.

18. Cannon, E.K., Portwood, J.L., 2nd, Hayford, R.K., Haley, O.C., Gardiner, J.M., Andorf, C.M. and Woodhouse, M.R. (2024) Enhanced pan-genomic resources at the maize genetics and genomics database. Genetics, 227.

19. Prasanna, B.M. (2012) Diversity in global maize germplasm: characterization and utilization. J Biosci, 37, 843–855.

20. Andorf, C., Beavis, W.D., Hufford, M., Smith, S., Suza, W.P., Wang, K., Woodhouse, M., Yu, J. and Lubberstedt, T. (2019) Technological advances in maize breeding: past, present and future. Theor Appl Genet, 132, 817–849.

21. Zhang, X., Liang, X. and Zhang, Y. (2025) Advancements in the Research and Application of Whole-Plant Maize Silage for Feeding Purposes. Animals (Basel), 15.

22. Kaur, G., Sethi, M., Devi, V., Kaur, A., Kaur, H. and Chaudhary, D.P. (2025) Investigating maize as a sustainable energy crop for bioethanol production: Delineating cultivation, utilization, biotechnological and environmental perspectives. Biomass Bioenergy, 198, 107867.

23. Cherwoo, L., Gupta, I., Flora, G., Verma, R., Kapil, M., Arya, S.K., Ravindran, B., Khoo, K.S., Bhatia, S.K., Chang, S.W., et al. (2023) Biofuels an alternative to traditional fossil fuels: A comprehensive review. Sustain. Energy Technol. Assessments, 60, 103503.

24. McCLINTOCK, B. (1950) The origin and behavior of mutable loci in maize. Proc Natl Acad Sci U S A, 36, 344–355.

25. Slotkin, R.K., Freeling, M. and Lisch, D. (2005) Heritable transposon silencing initiated by a naturally occurring transposon inverted duplication. Nat Genet, 37, 641–644.

26. Stitzer, M.C., Anderson, S.N., Springer, N.M. and Ross-Ibarra, J. (2021) The genomic ecosystem of transposable elements in maize. PLoS Genet, 17, e1009768.

27. Creighton, H.B. and McClintock, B. (1931) A Correlation of Cytological and Genetical Crossing-Over in Zea Mays. Proc Natl Acad Sci U S A, 17, 492–497.

28. Wei, F., Coe, E., Nelson, W., Bharti, A.K., Engler, F., Butler, E., Kim, H., Goicoechea, J.L., Chen, M., Lee, S., et al. (2007) Physical and genetic structure of the maize genome reflects its complex evolutionary history. PLoS Genet, 3, e123.

29. Wang, B., Hou, M., Shi, J., Ku, L., Song, W., Li, C., Ning, Q., Li, X., Li, C., Zhao, B., et al. (2023) De novo genome assembly and analyses of 12 founder inbred lines provide insights into maize heterosis. Nat Genet, 55, 312–323.

30. Hufford, M.B., Seetharam, A.S., Woodhouse, M.R., Chougule, K.M., Ou, S., Liu, J., Ricci, W.A., Guo, T., Olson, A., Qiu, Y., et al. (2021) De novo assembly, annotation, and comparative analysis of 26 diverse maize genomes. Science, 373, 655–662.

31. Stitzer, M., Seetharam, A., Armin, S., Hsu, S.-K., Schulz, A., Taylor, A.-E., Hale, C., Syring, M., Minx, P., Pasquet, R., et al. (2024) Extensive genome evolution distinguishes maize within a stable tribe of grasses. 10.5281/ZENODO.10683652.

32. Sen, T.Z., Harper, L.C., Schaeffer, M.L., Andorf, C.M., Seigfried, T.E., Campbell, D.A. and Lawrence, C.J. (2010) Choosing a genome browser for a Model Organism Database: surveying the maize community. Database (Oxford), 2010, baq007.

33. Woodhouse, M.R., Cannon, E.K., Portwood, J.L., 2nd, Harper, L.C., Gardiner, J.M., Schaeffer, M.L. and Andorf, C.M. (2021) A pan-genomic approach to genome databases using maize as a model system. BMC Plant Biol, 21, 385.

34. Woodhouse, M.R., Portwood, J.L., Sen, S., Hayford, R.K., Gardiner, J.M., Cannon, E.K., Harper, L.C. and Andorf, C.M. (2023) Maize protein structure resources at the maize genetics and genomics database. Genetics, 224.

35. Cao, Y., Dou, D., Zhang, D., Zheng, Y., Ren, Z., Su, H., Sun, C., Hu, X., Bao, M., Zhu, B., et al. (2022) ZmDWF1 regulates leaf angle in maize. Plant Sci, 325, 111459.

36. Sarkar, B., Varalaxmi, Y., Vanaja, M., RaviKumar, N., Prabhakar, M., Yadav, S.K., Maheswari, M. and Singh, V.K. (2023) Mapping of QTLs for morphophysiological and yield traits under water-deficit stress and well-watered conditions in maize. Front Plant Sci, 14, 1124619.

37. Sharma, J., Sharma, S., Sai Karnatam, K., Prakash Raigar, O., Lahkar, C., Kumar Saini, D., Kumar, S., Singh, A., Kumar Das, A., Sharma, P., et al. (2023) Surveying the genomic landscape of silage-quality traits in maize (Zea mays L.). Crop J., 11, 1893–1901.

38. Wallace, J.G., Bradbury, P.J., Zhang, N., Gibon, Y., Stitt, M. and Buckler, E.S. (2014) Association mapping across numerous traits reveals patterns of functional variation in maize. PLoS Genet, 10, e1004845.

39. Song, B., Marco-Sola, S., Moreto, M., Johnson, L., Buckler, E.S. and Stitzer, M.C. (2022) AnchorWave: Sensitive alignment of genomes with high sequence diversity, extensive structural polymorphism, and whole-genome duplication. Proc Natl Acad Sci U S A, 119.

40. Wei, W., Gui, S., Yang, J., Garrison, E., Yan, J. and Liu, H.-J. (2025) wgatools: an ultrafast toolkit for manipulating whole-genome alignments. Bioinformatics, 41.

41. Guarracino, A., Heumos, S., Nahnsen, S., Prins, P. and Garrison, E. (2022) ODGI: understanding pangenome graphs. Bioinformatics, 38, 3319–3326.

42. Garrison, E., Siren, J., Novak, A.M., Hickey, G., Eizenga, J.M., Dawson, E.T., Jones, W., Garg, S., Markello, C., Lin, M.F., et al. (2018) Variation graph toolkit improves read mapping by representing genetic variation in the reference. Nat Biotechnol, 36, 875–879.

43. Gruning, B., Dale, R., Sjodin, A., Chapman, B.A., Rowe, J., Tomkins-Tinch, C.H., Valieris, R., Koster, J. and Bioconda Team (2018) Bioconda: sustainable and comprehensive software distribution for the life sciences. Nat Methods, 15, 475–476.

44. Wickham, H. (2016) Ggplot2: Elegant graphics for data analysis 2nd ed. Springer International Publishing, Cham, Switzerland.

45. Schulz, A.J., Zhai, J., AuBuchon-Elder, T., Andorf, C.M., El-Walid, M.Z., Ferebee, T.H., Gilmore, E.H., Hufford, M.B., Johnson, L.C., Kellogg, E.A., et al. (2025) Fishing for a reelGene: evaluating gene models with evolution and machine learning. Plant J, 123, e70483.

46. van Wijk, K., Leppert, T., Sun, Z., Guzchenko, I., Debley, E., Sauermann, G., Routray, P., Mendoza, L., Sun, Q. and Deutsch, E. (2023) TheZea maysPeptideAtlas - a new maize community resource. bioRxiv, 10.1101/2023.12.21.572651.

47. Cagirici, H.B., Budak, H. and Sen, T.Z. (2022) G4Boost: a machine learning-based tool for quadruplex identification and stability prediction. BMC Bioinformatics, 23, 240.

48. Piazza, A., Boule, J.-B., Lopes, J., Mingo, K., Largy, E., Teulade-Fichou, M.-P. and Nicolas, A. (2010) Genetic instability triggered by G-quadruplex interacting Phen-DC compounds in Saccharomyces cerevisiae. Nucleic Acids Res, 38, 4337–4348.

49. Yadav, P., Harcy, V., Argueso, J.L., Dominska, M., Jinks-Robertson, S. and Kim, N. (2014) Topoisomerase I plays a critical role in suppressing genome instability at a highly transcribed G-quadruplex-forming sequence. PLoS Genet, 10, e1004839.

50. Lexa, M., Kejnovsky, E., Steflova, P., Konvalinova, H., Vorlfckova, M. and Vyskot, B. (2014) Quadruplex-forming sequences occupy discrete regions inside plant LTR retrotransposons. Nucleic Acids Res, 42, 968–978.

51. Andorf, C., Ross-Ibarra, J., Seetharam, A., Hufford, M. and Woodhouse, M. (2024) A unified VCF data set from nearly 1,500 diverse maize accessions and resources to explore the genomic landscape of maize. bioRxiv, 10.1101/2024.04.30.591904.

52. Cannon, S.B., Lee, H.-O., Weeks, N.T. and Berendzen, J. (2024) Pandagma: a tool for identifying pan-gene sets and gene families at desired evolutionary depths and accommodating whole-genome duplications. Bioinformatics, 40.

53. Haas, B.J., Delcher, A.L., Wortman, J.R. and Salzberg, S.L. (2004) DAGchainer: a tool for mining segmental genome duplications and synteny. Bioinformatics, 20, 3643–3646.

